# Menopause, Brain Anatomy, Cognition and Alzheimer’s Disease

**DOI:** 10.1101/2022.10.18.512730

**Authors:** Manuela Costantino, Grace Pigeau, Olivier Parent, Justine Ziolkowski, Gabriel A. Devenyi, Nicole J. Gervais, M. Mallar Chakravarty

## Abstract

The menopause transition has been repeatedly associated with decreased cognitive performance and increased incidence of Alzheimer’s Disease (AD), particularly when it is induced surgically ^1,2^ or takes place at a younger age ^3,4^. However, there are very few studies that use neuroimaging techniques to examine the effects of these variables in aggregate and in a large sample. Here, we use data from thousands of participants from the UK Biobank to assess the relationship between menopausal status, menopause type (surgical or natural), and age at menopause with cognition, AD, and neuroanatomical measures derived from magnetic resonance imaging. We find that for brain and cognitive measures, menopausal status, menopause type and age at surgical menopause do not impact the brain; but that there is a positive correlation between anatomy, cognition and age at non-surgical menopause. These results do not align with previous reports in the literature with smaller samples. However, we confirm that both early and surgical menopause are associated with a higher risk of developing AD, indicating that early and abrupt ovarian hormone deprivation might contribute to the development of the disorder.

## Introduction

Menopause is associated with a decrease in circulating ovarian hormones, which impacts multiple physiological systems, including brain structure and function ^5,6^. Surgical menopause (where ovaries are removed via bilateral oophorectomy) as opposed to spontaneous (after one year without menstruation) leads to a faster decrease in hormone production. The timing of the transition is also believed to impact women’s health, thus, surgical and early menopause have been used to model the impact that ovarian hormone deprivation may have on the ageing process. Critically, lower cognitive performance and increased incidence of Alzheimer’s Disease (AD), have been observed in women who had their ovaries removed prior to non-surgical menopause ^7^. Similarly, early non-surgical menopause has been associated with accelerated neurodegeneration, measured with magnetic resonance imaging (MRI), in addition to worse cognition ^3,4^. Adding to this critical body of literature are mixed findings related to improved cognition in women taking hormone therapy (HT), exogenous female hormones, during the transition ^8^. Although these vary in type, dose, duration and timing, they contain estrogens with or without progestin ^2^. Thus, we investigated how both surgical and spontaneous menopause, as well as age at menopause, related to multiple measures of cognitive performance, brain anatomy, and AD incidence in a large cohort of women from the UK Biobank, while robustly controlling for age.

## Methods

### Sample

Cross-sectional demographic, cognitive, MRI and health-outcome data was obtained from the UK Biobank. Based on self-report data, female participants were divided into premenopausal women (PRE), women who underwent a bilateral oophorectomy (with or without a hysterectomy) prior to menopause (SURG) and women who underwent menopause without surgical interference (POST). Participants who reported uncertainty of their menopausal status or who underwent a hysterectomy without a bilateral oophorectomy were excluded from the study. Different subsamples of these groups were included in each analysis, based on data availability (Table 1).

**Table 1:**
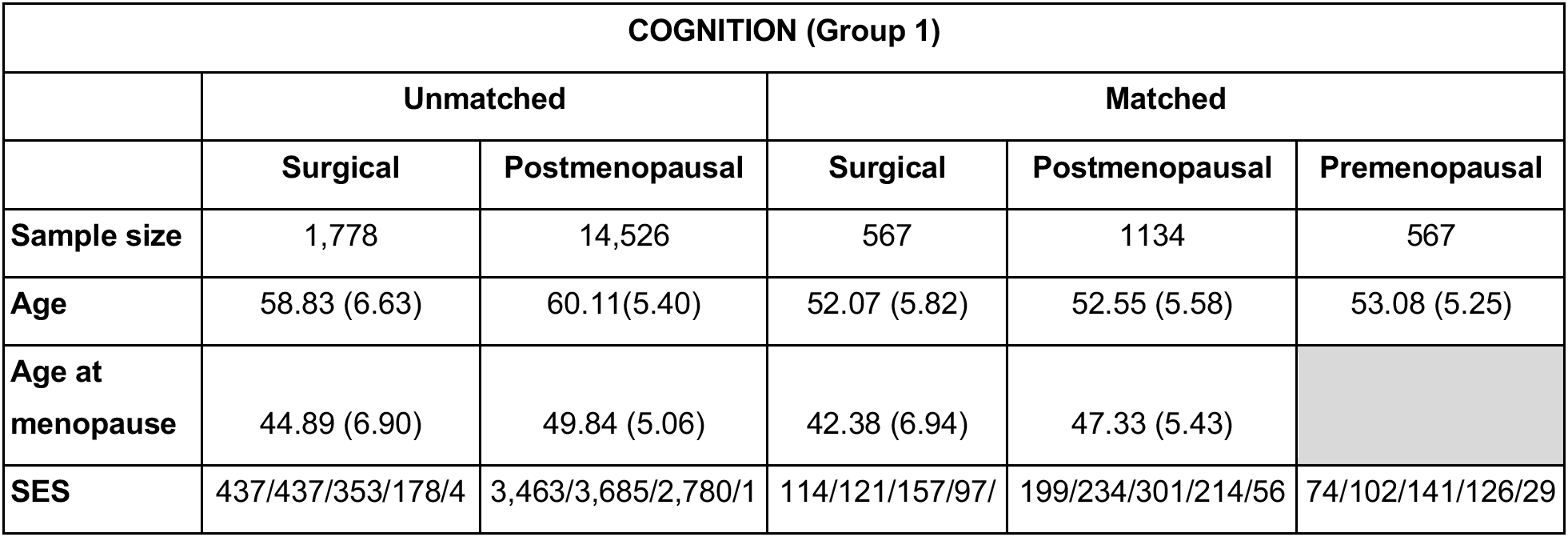
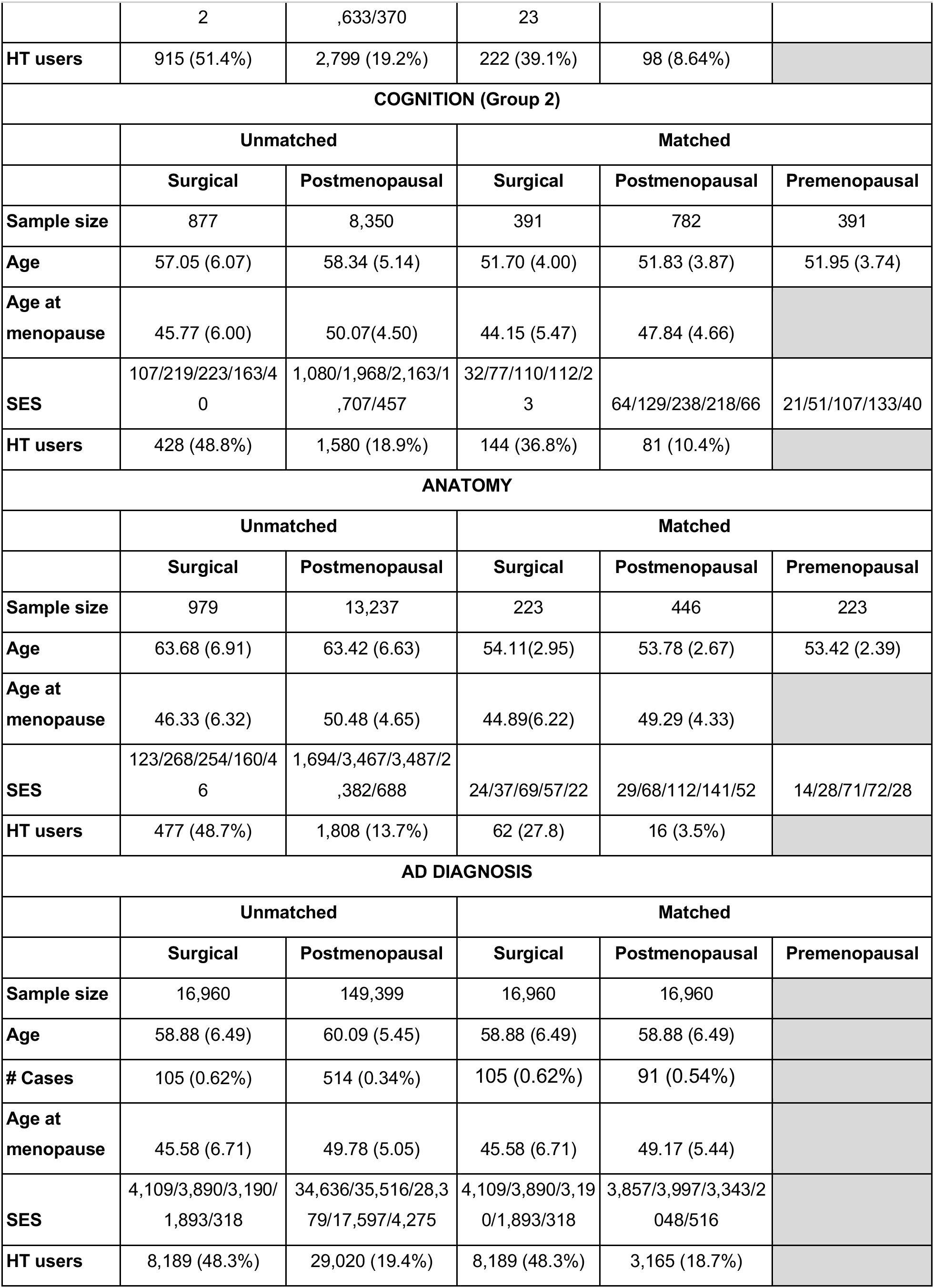
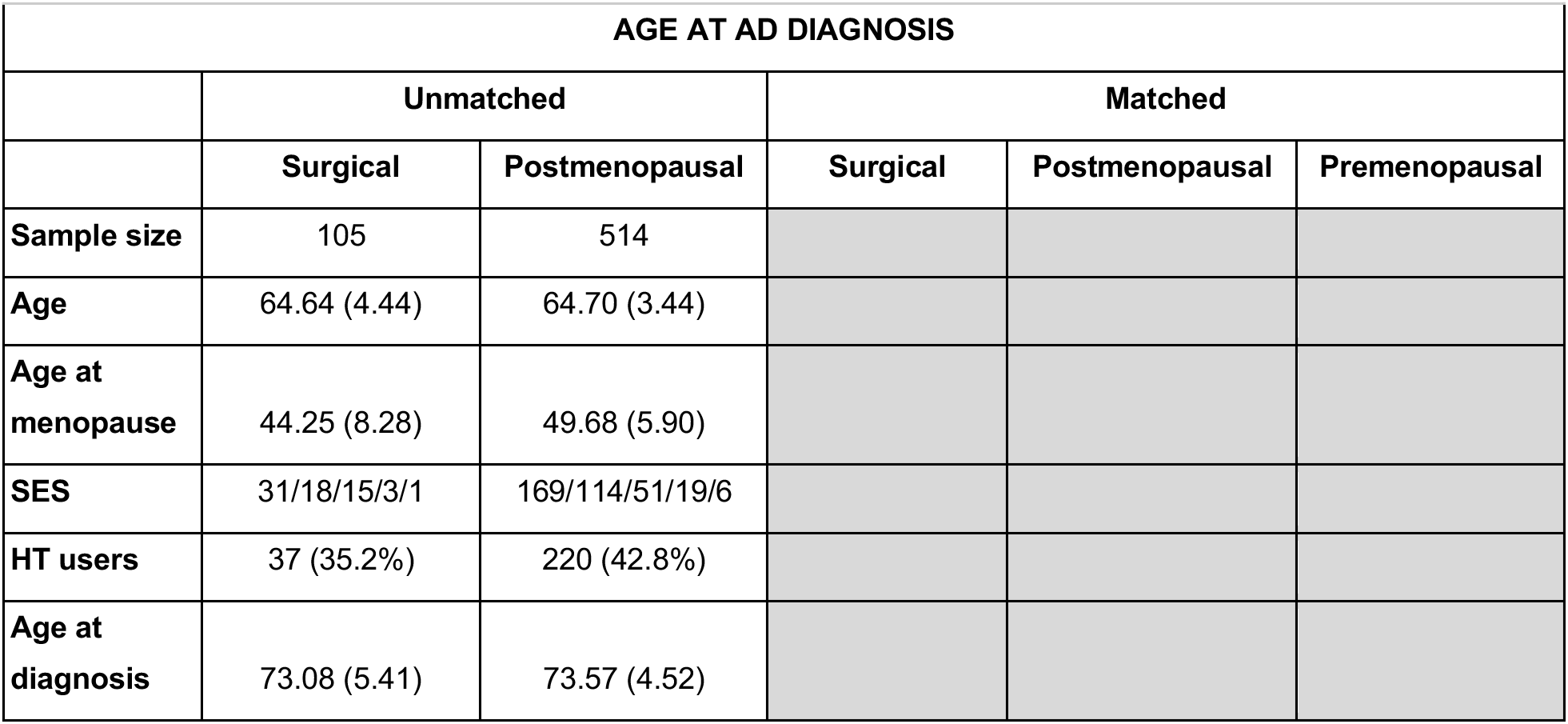
Basic demographic information for the participants in each analysis. For age and age at menopause, the mean and standard deviation, in parentheses, are shown. The number of participants who used HT as well as the proportion of the total sample that they represent is also reported. For SES, the values represent the number of participants in each of the following income categories, in pounds sterling: less than 18,000/18,000-30,999/31,000-51,999/52,000-100,000/greater than 100,000. Over 90% of the sample identifies as ethnically white.

| COGNITION (Group 1) |  |  |  |  |  |
| --- | --- | --- | --- | --- | --- |
|  | Unmatched |  | Matched |  |  |
|  | Surgical | Postmenopausal | Surgical | Postmenopausal | Premenopausal |
| **Sample size** | 1,778 | 14,526 | 567 | 1134 | 567 |
| **Age** | 58.83 (6.63) | 60.11(5.40) | 52.07 (5.82) | 52.55 (5.58) | 53.08 (5.25) |
| **Age at menopause** | 44.89 (6.90) | 49.84 (5.06) | 42.38 (6.94) | 47.33 (5.43) |  |
| **SES** | 437/437/353/178/4 | 3,463/3,685/2,780/1 | 114/121/157/97/ | 199/234/301/214/56 | 74/102/141/126/29 |
|  | 2 | ,633/370 | 23 |  |  |
| **HT users** | 915 (51.4%) | 2,799 (19.2%) | 222 (39.1%) | 98 (8.64%) |  |
| **COGNITION (Group 2)** |  |  |  |  |  |
|  | **Unmatched** |  | **Matched** |  |  |
|  | **Surgical** | **Postmenopausal** | **Surgical** | **Postmenopausal** | **Premenopausal** |
| **Sample size** | 877 | 8,350 | 391 | 782 | 391 |
| **Age** | 57.05 (6.07) | 58.34 (5.14) | 51.70 (4.00) | 51.83 (3.87) | 51.95 (3.74) |
| **Age at menopause** | 45.77 (6.00) | 50.07(4.50) | 44.15 (5.47) | 47.84 (4.66) |  |
| **SES** | 107/219/223/163/4<br>0 | 1,080/1,968/2,163/1<br>,707/457 | 32/77/110/112/2<br>3 | 64/129/238/218/66 | 21/51/107/133/40 |
| **HT users** | 428 (48.8%) | 1,580 (18.9%) | 144 (36.8%) | 81 (10.4%) |  |
| **ANATOMY** |  |  |  |  |  |
|  | **Unmatched** |  | **Matched** |  |  |
|  | **Surgical** | **Postmenopausal** | **Surgical** | **Postmenopausal** | **Premenopausal** |
| **Sample size** | 979 | 13,237 | 223 | 446 | 223 |
| **Age** | 63.68 (6.91) | 63.42 (6.63) | 54.11(2.95) | 53.78 (2.67) | 53.42 (2.39) |
| **Age at menopause** | 46.33 (6.32) | 50.48 (4.65) | 44.89(6.22) | 49.29 (4.33) |  |
| **SES** | 123/268/254/160/4<br>6 | 1,694/3,467/3,487/2<br>,382/688 | 24/37/69/57/22 | 29/68/112/141/52 | 14/28/71/72/28 |
| **HT users** | 477 (48.7%) | 1,808 (13.7%) | 62 (27.8) | 16 (3.5%) |  |
| **AD DIAGNOSIS** |  |  |  |  |  |
|  | **Unmatched** |  | **Matched** |  |  |
|  | **Surgical** | **Postmenopausal** | **Surgical** | **Postmenopausal** | **Premenopausal** |
| **Sample size** | 16,960 | 149,399 | 16,960 | 16,960 |  |
| **Age** | 58.88 (6.49) | 60.09 (5.45) | 58.88 (6.49) | 58.88 (6.49) |  |
| **# Cases** | 105 (0.62%) | 514 (0.34%) | 105 (0.62%) | 91 (0.54%) |  |
| **Age at menopause** | 45.58 (6.71) | 49.78 (5.05) | 45.58 (6.71) | 49.17 (5.44) |  |
| **SES** | 4,109/3,890/3,190/<br>1,893/318 | 34,636/35,516/28,3<br>79/17,597/4,275 | 4,109/3,890/3,19<br>0/1,893/318 | 3,857/3,997/3,343/2<br>048/516 |  |
| **HT users** | 8,189 (48.3%) | 29,020 (19.4%) | 8,189 (48.3%) | 3,165 (18.7%) |  |

**Table 1:** Basic demographic information for the participants in each analysis.
| AGE AT AD DIAGNOSIS |  |  |  |  |  |
| --- | --- | --- | --- | --- | --- |
|  | Unmatched |  | Matched |  |  |
|  | Surgical | Postmenopausal | Surgical | Postmenopausal | Premenopausal |
| **Sample size** | 105 | 514 |  |  |  |
| **Age** | 64.64 (4.44) | 64.70 (3.44) |  |  |  |
| **Age at menopause** | 44.25 (8.28) | 49.68 (5.90) |  |  |  |
| **SES** | 31/18/15/3/1 | 169/114/51/19/6 |  |  |  |
| **HT users** | 37 (35.2%) | 220 (42.8%) |  |  |  |
| **Age at diagnosis** | 73.08 (5.41) | 73.57 (4.52) |  |  |  |

Participants were excluded if self-reported data was missing for menopausal status or type, age at menopause or use of HT. Socioeconomic status (SES) was quantified based on total household income and missing data was imputed by the mean. HT was encoded as a binary variable, distinguishing those with a history of HT during the menopause transition and those without it. Unfortunately, no additional information about HT was available in UK Biobank records. Date of AD diagnosis was obtained from hospital admissions data linked to the dataset. ^9^

### Cognitive Data

Cognitive tests were performed at the UK Biobank’s assessment centers ^10^. Seven domains were selected, based on data availability: fluid intelligence score, numeric memory, pairs matching, prospective memory, reaction time, symbol digit substitution and tower rearrangement. To maximize power while avoiding missing data, participants were included if they had participated in all of the first five tests (cognitive group 1) or in both of the last two (cognitive group 2) (Table 1).

### MRI Data

T1-weighted structural MRI scans were processed with CIVET 2.1.1 and four morphometric outputs were selected: total brain volume (TBV), total gray matter volume (GMV), total white matter volume (WMV) and vertex-wise cortical thickness (CT) measurements at 77,122 points across the cortical surface. Participants were further excluded based on manual quality control of raw scans and cortical reconstructions.

### Statistical Analysis

Groups were age matched using the nearest neighbors algorithm with the MatchIt package in R 4.1.2 and the supplementary methods can be found in the online-only materials. Multiple regressions were performed with the seven cognitive tests and the three whole-brain anatomical measures as response variables. For each of them, three models were used:

*Model 1: (cognitive/anatomical measure) ~ age + group + SES*
*Model 2: (cognitive/anatomical measure) ~ age + age at menopause + HT + SES, POST only*
*Model 3: (cognitive/anatomical measure) ~ age + age at menopause + HT + SES, SURG only*

Multiple comparisons were accounted for using Bonferroni correction. These same three models were carried out on a vertex-wise basis, with CT as a response variable and false detection rate (FDR) correction. A chi squared test was performed to compare the prevalence of AD in the POST and SURG groups and models 2 and 3 were used to examine the effects of age at menopause on age at AD diagnosis. Logistic regressions with the same predictors as models 2 and 3 were performed with AD diagnosis as a response variable. Analyses were run in R v4.1.2 and using RMINC v.1.5.3.0.

### Supplementary analyses

Alternative models were attempted, including the encoding of age at menopause as a categorical variable (premature: <40, early: <45, spontaneous: 45+) (Supplementary fig 1), the inclusion of a group by age at menopause interaction term (Supplementary fig 2), the removal of all participants under the age of 60 (Supplementary fig 3) and the removal of all participants who with a history of HT (Supplementary fig 4)

## Results

A full description of the samples can be found in Table 1. 3,373 MRIs were discarded after quality control, namely 151(14.7%) in the PRE group, 3,091(18.9%) in the POST group and 313(23.8%) in the SURG group.

### Anatomy and Cognition

We observed that later onset of menopause in the POST group only was positively associated with fluid intelligence (*β*=0.095, p<0.001), numeric memory (*β*=0.064, p<0.001) and pairs matching (*β*=0.040, p<0.001). These observations are further supported by results that demonstrate total brain (*β*=0.046, p<0.001), white matter (*β*=0.047, p<0.001), and grey matter volumes (*β*=0.033, p<0.001) being relatively preserved in those with later onset of menopause in the POST group (see Fig 1a). All the continuous variables were z-scored to obtain standardized beta coefficients. Further still, age at non-surgical menopause was positively associated with CT in many regions across the cortical surface (Fig 1b). Outside of these findings age of menopause, age and HT status were not associated with cognitive performance or brain anatomy measures.

**Figure:**
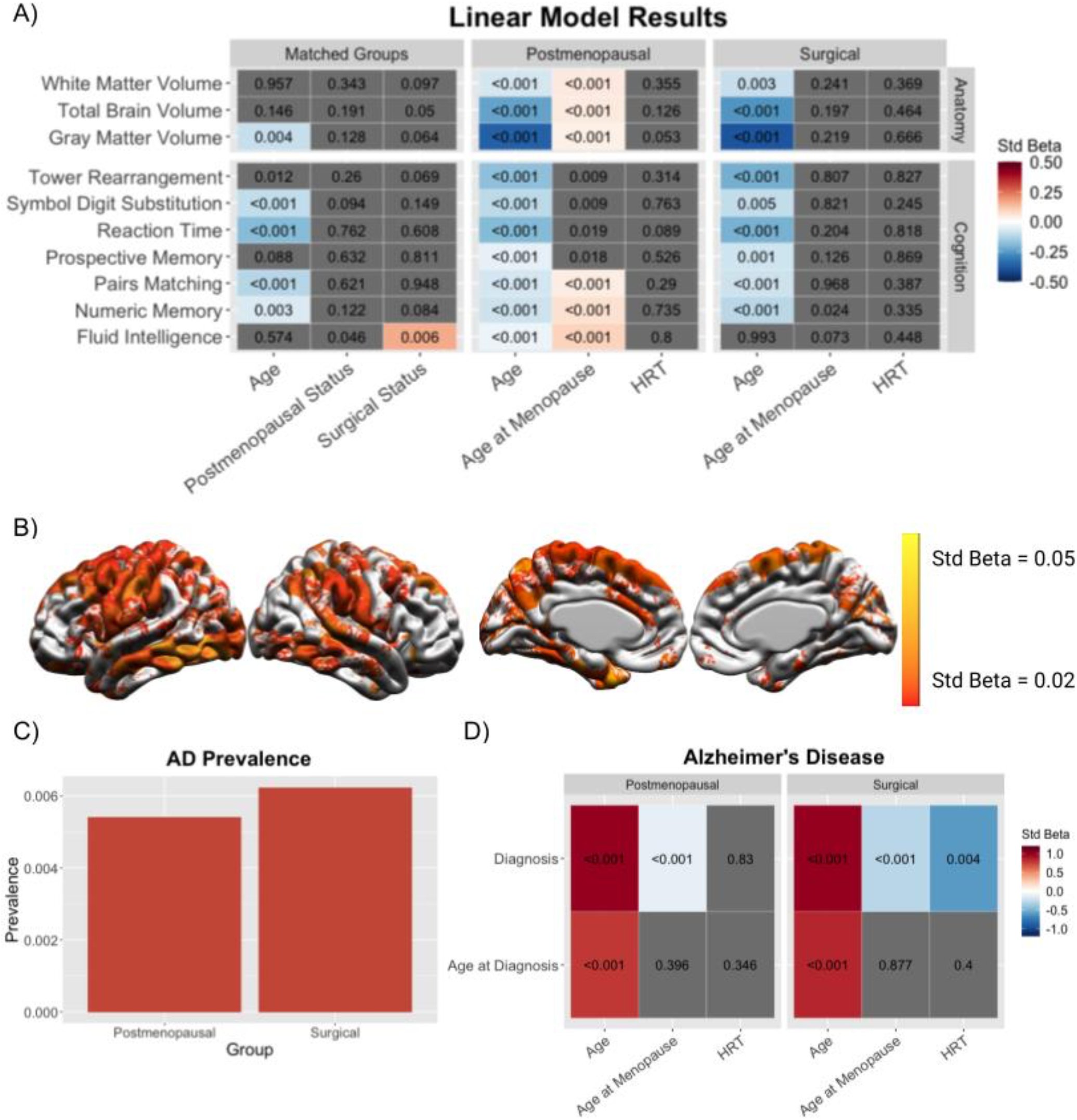
A) Results from multiple regressions - Each row within each panel represents a linear model with three predictors {bottom) and a response var able (left). The number w thin each cell s the uncorrected p-value for the predictor, rounded to 3 decimal places, and cells are colored in grey when the corrected p value is larger than 0 05 Colors represent the standardized beta coeff cients of the significant predictors. SES was corrected for in every model. B) Standardized beta coeff cients of age at non surgical menopause in a vertex-w se linear model w th cortical thickness as a response variable. Colored vert ces are significant at 1% FOR. C) Relat ve prevalence of A zheimer s disease in women who underwent surgical and non-surgical menopause. D) Results of the logistic regressions with AD diagnosis as a response var able and of the multip e regress ons with age at AD diagnosis as a response variable. The plot can be interpreted in the same way as figure A and SES was also ncluded in all models.

### Alzheimer’s Disease

The rates of conversion to AD (see Fig 1c and d) further supports a cognitive and brain preservation effect of a later onset of menopause. Logistic and multiple regressions and chi squared tests suggest that women in the SURG group are more likely to have AD (**χ**2 = 1.0058, p = 0.3159) and that an earlier age of menopause is related to higher rates of AD diagnosis (*β*=−0.1091, p<0.001 for POST, *β*=−0.2879, p<0.001 for SURG). In the SURG group, women who did not consume HT are more likely to develop AD (*β*=− 0.6003, p=0.004).

## Discussion

Overall, our findings suggest that menopause status does not have notable effects on the brain or cognition, for both surgical and non-surgical groups. These critical findings do not agree with previous reports of brain atrophy and reduced cognition ^11,12^, particularly for those undergoing surgical menopause ^1,13^. This highlights the importance of larger sample sizes in association studies to prevent incidental findings. Nontheless, our findings suggest that those undergoing non-surgical menopause at an earlier age show domain-specific cognitive decline and brain atrophy patterns, as previously reported ^3,14^. Spontaneous, but not surgical, menopause depends on multiple factors such as genetics, lifestyle factors and comorbidities ^15^, which could explain this difference and may be intertwined as causal factors that impact this relationship. Overall, our results underscore the urgent need to increase sample sizes and improve age-correction as a means of understanding sex-specific factors that may underly dementia.

Our findings of fewer AD diagnoses in those with a later menopause onset further support its protective effect in women with surgical menopause, suggesting that early and abrupt hormone deprivation predisposes women to AD later in life without impacting their cognition or brain health at midlife ^16^. Any of the endocrine, immune, or cardiovascular changes associated with menopause could be interacting with AD patopathology to alter its progression, as opposed to directly worsening general brain health.

The question of whether the relationship between menopause and the brain’s structural and functional changes is causal remains unanswered. On the one hand, there is evidence from animal studies that ovarian hormone deprivation causes changes in the brain and in behavior and there are multiple proposed pathways ^17^. Yet, our results do not show an impact of menopausal status and the positive correlation between age at menopause and brain health might simply be explained by generally healthy women having both healthier brains and later menopause. In either case, our study confirms that there are relevant interactions between the endocrine and nervous systems, which supports the idea that brain aging and neurodegenerative diseases should not be studied in isolation, but rather by integrating data from multiple physiological systems.

Some limitations to the study are the lack of a perimenopausal group and of distinction between HT regimens, due to data availability. Most demographic and reproductive variables were self-reported, which is known to introduce noise and bias, such as incorrect determination of menopausal status. Furthermore, we do not have any measure of verbal memory, which is known to decrease during perimenopause ^18^. Finally, the proportion of the cohort that was diagnosed with AD is limited, particularly in the SURG group. Therefore, it will be critical to monitor the AD-conversion rates as UK Biobank continues to collect data. Since this study only examines the long term effects of the transition, no conclusions can be drawn on how perimenopause affects the brain.

Thus, our study is the first to comprehensively analyze well-powered MRI, cognitive, and clinical data in the context of menopause with robust age correction. We observed that, in healthy populations, menopause was not associated with changes in the brain, except in women who underwent early spontaneous menopause. However, both surgical transitions and early age at menopause of any type increased the risk of AD. Further research is needed to understand the mechanisms behind the interactions between these two systems.

## Notes

### Competing Interest Statement

The authors have declared no competing interest.

