## Supplementary Materials for "Menopause, Brain Anatomy, Cognition and Alzheimer’s Disease"

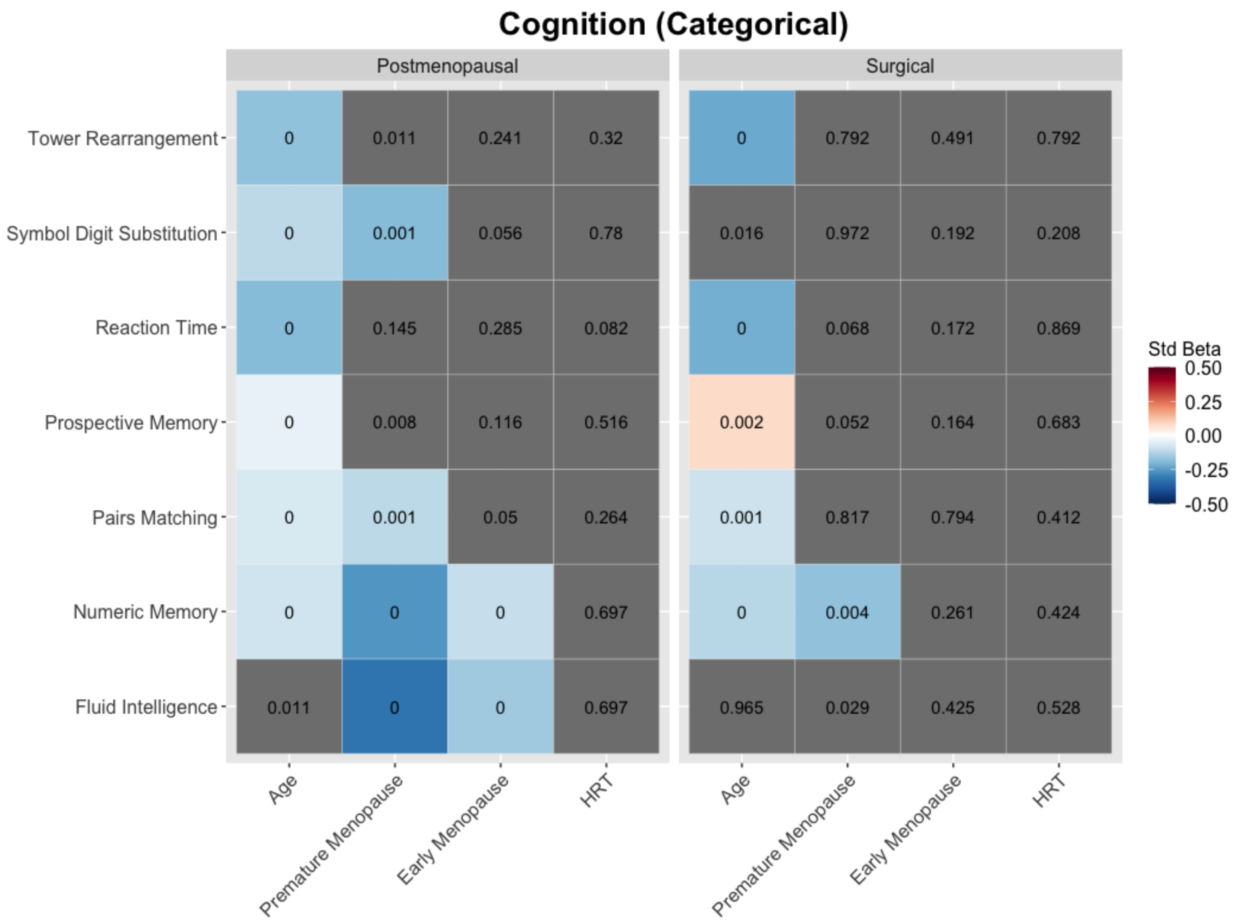


**Supplementary Figure 1**: Multiple regressions examining the effect of age, age at menopause and HRT on the 7 cognitive variables. Instead of using it as a continuous variable, age at menopause was encoded as a three level factor: premature ( <40 years), early (<45) and spontaneous (45+), which was set as the reference level. Gray cells represent predictors that were not found to be significant after multiple comparisons correction.

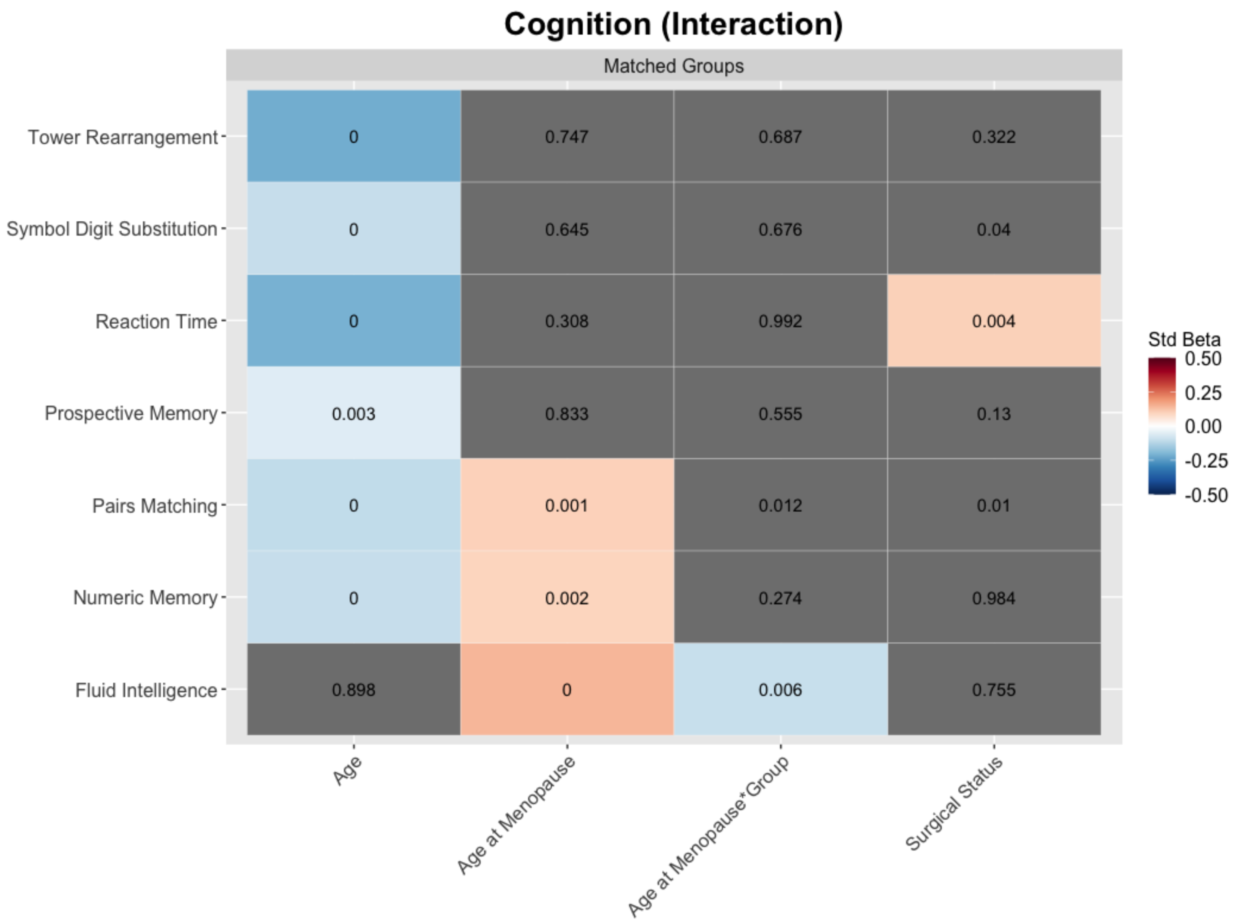


**Supplementary Figure 2**: Multiple regressions examining the effect of age, age at menopause, menopause type and a age at menopause by menopause type interaction term on age-matched women of the POST and SURG groups.

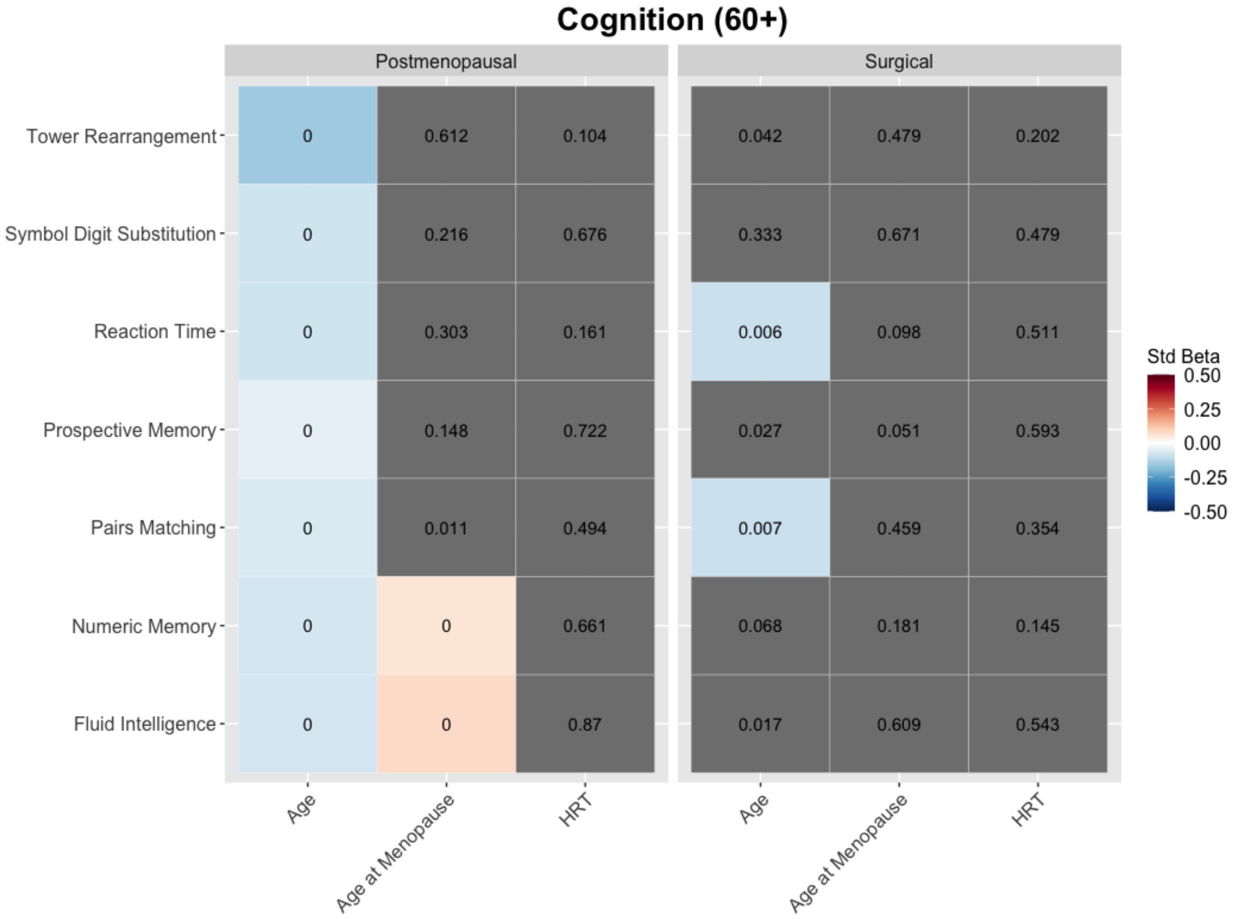


**Supplementary Figure 3**: Multiple regressions examining the effect of age, age at menopause and HRT on cognition in women who were over the age of 60 at the time of assessment

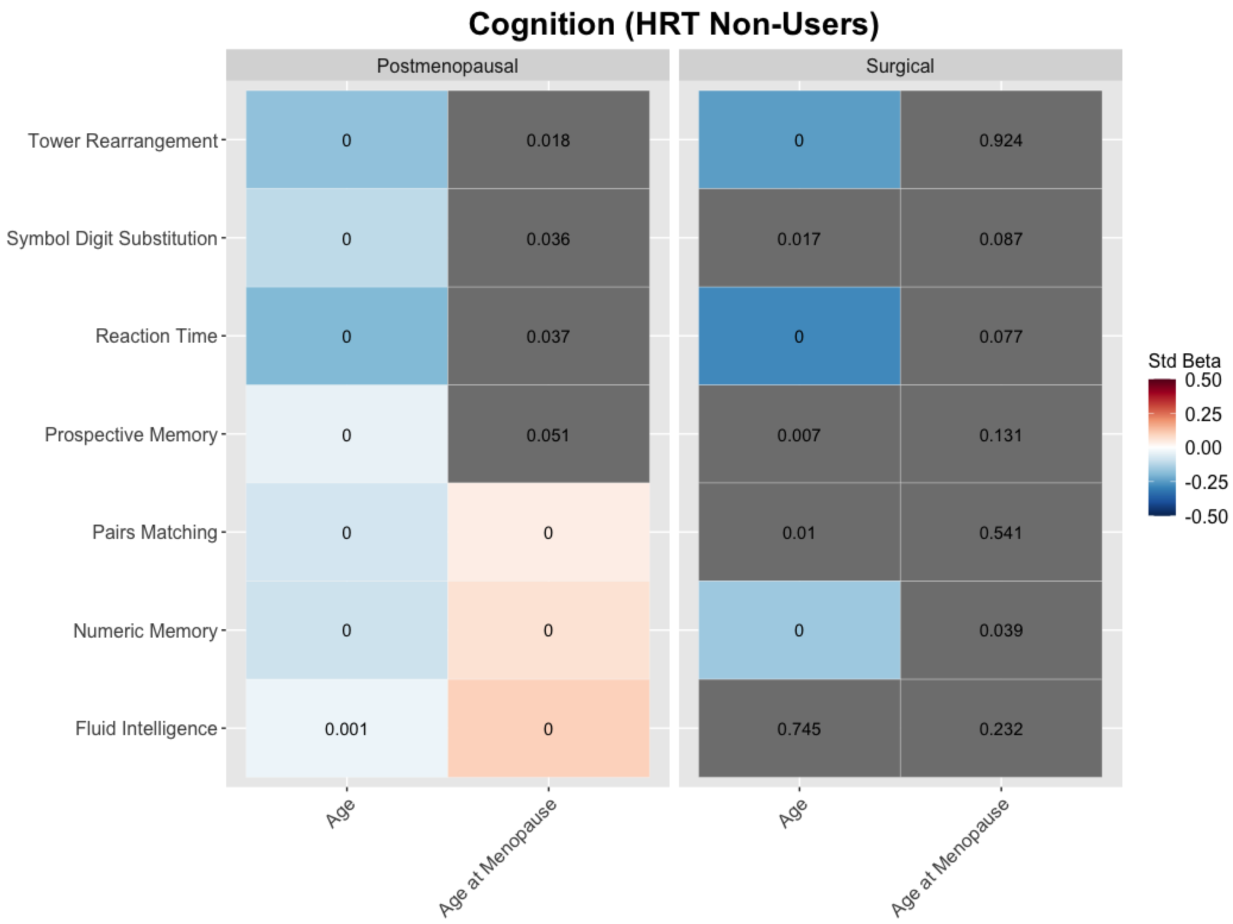


**Supplementary Figure 4**: Multiple regressions examining the effect of age and age at menopause on cognition in women who did not consume HRT during the menopause transition.

**Supplementary Methods**: For analyses of neuroanatomy and cognition, PRE and SURG groups were age-matched using the k-nearest neighbors algorithm. Distance was estimated with propensity scores using the MatchIt v4.3.4 package in R 4.1.2. The POST group was then matched to the resulting participants using the same method. For the analysis on AD prevalence, the SURG and POST groups were directly matched.
